# Obsessive-Compulsive Disorder (OCD) is Associated with Increased Electroencephalographic (EEG) Delta and Theta Oscillatory Power but Reduced Delta Connectivity

**DOI:** 10.1101/2022.10.03.510571

**Authors:** M. Prabhavi N. Perera, Sudaraka Mallawaarachchi, Neil W. Bailey, Oscar W. Murphy, Paul B. Fitzgerald

**Affiliations:** Central Clinical School, Monash University, Wellington Road, Clayton, Victoria 3800, Australia; Melbourne Integrative Genomics, School of Mathematics & Statistics, University of Melbourne, Parkville, Victoria 3052, Australia; Bionics Institute, East Melbourne, Victoria 3002, Australia; School of Medicine and Psychology, Australian National University, Canberra, ACT 2600, Australia

**Keywords:** Cluster-Based Permutation, Electroencephalography, Functional Connectivity, Obsessive-Compulsive Disorder, Power Spectral Analysis, 1/f non-oscillatory activity

## Abstract

**Background:** Obsessive-Compulsive Disorder (OCD) is a mental health condition causing significant decline in the quality of life of sufferers and the limited knowledge on the pathophysiology hinders successful treatment. The aim of the current study was to examine electroencephalographic (EEG) findings of OCD to broaden our understanding of the disease.

**Methods:** Resting-state eyes-closed EEG data was recorded from 25 individuals with OCD and 27 healthy controls (HC). The 1/f arrhythmic activity was removed prior to computing oscillatory powers of all frequency bands (delta, theta, alpha, beta, gamma). Cluster-based permutation was used for between-group statistical analyses, and comparisons were performed for the 1/f slope and intercept parameters. Functional connectivity (FC) was measured using coherence and debiased weighted phase lag index (d-wPLI), and statistically analysed using the Network Based Statistic method.

**Results:** Compared to HC, the OCD group showed increased oscillatory power in the delta and theta bands in the fronto-temporal and parietal brain regions. However, there were no significant between-group findings in other bands or 1/f parameters. The coherence measure showed significantly reduced FC in the delta band in OCD compared to HC but the d-wPLI analysis showed no significant differences.

**Conclusions:** OCD is associated with raised oscillatory power in slow frequency bands in the fronto-temporal brain regions, which agrees with the previous literature and therefore is a potential biomarker. Although delta coherence was found to be lower in OCD, due to inconsistencies found between measures and the previous literature, further research is required to ascertain definitive conclusions.

## 1. INTRODUCTION

Obsessive-compulsive disorder (OCD) is a mental health disorder characterized by unwanted and intrusive thoughts (obsessions), often leading to compulsions such as repetitive behaviors, mental acts, or rigidly applied rituals [1]. With a lifetime prevalence of 1-3%, OCD has been classified as a mental health condition causing significant reduction in the quality of life of sufferers [2]. First-line treatments for OCD, including Selective Serotonin Reuptake Inhibitors (SSRIs) and cognitive behavioral therapy are found to have poor success rates [3]. This inadequacy has motivated investigators to study the underlying pathophysiology of OCD in the hope of identifying better suited therapeutic approaches.

Resting-state electroencephalographic (EEG) studies have shown that individuals with OCD show significant differences in brain electrical activity compared to healthy controls (HC) [4]. Although differences in the power of neural oscillations within all frequency bands have been reported in the previous literature, reports of delta, theta and alpha differences are the most common. Several studies have noted increased resting fronto-temporal delta [5, 6] and theta [7, 8] power in OCD groups compared to HC. However, there are inconsistencies between the studies examining alpha activity, with reports of both power increases [9, 10] and decreases [11, 12].

Identification and analysis of brain oscillatory activity involves transforming the EEG data to frequency or time-frequency domain. Power in the frequency spectrum reflects both oscillatory activity and irregular, asynchronous, aperiodic firing of neurons or assemblies of neurons and exhibits a 1/f-like power spectrum, where power (amplitude) decreases as a function of frequency [13]. When measuring oscillatory power, it is imperative to correct for this non-oscillatory activity to avoid confounding estimates of power, obscuring differences in oscillatory activity between bands, and creating false differences where none exist. Therefore, detecting the rhythmicity of brain oscillations needs to be an additional consideration when interpreting measures of oscillatory power. Additionally, it is important to perform between-group analyses of the 1/f slope and intercept parameters as these may better represent the neural activity at lower frequencies and may also be functionally and behaviorally relevant [14].

Functional connectivity (FC) refers to various measures of how neural activity of one brain region relates to activity in another [15]. Several studies have reported altered connectivity in OCD when compared to HC. Many of these studies have reported decreased connectivity in alpha and beta frequency bands [16–20], while one study reported decreased global field synchronization of frontal EEG in the delta band [21]. Furthermore, it has been hypothesized that the pathophysiology of OCD may be attributed to defective structure and function of the frontal-striatal-thalamic (FST) circuit, resulting in poor FC between these regions [22]. However, the substantial heterogeneity between data collection parameters, EEG pre-processing methods and the utilized connectivity measures makes it difficult to interpret these results.

A principal aim of the current study was to investigate the differences in EEG power spectra in all frequency bands between OCD and HC groups. The primary hypothesis was that individuals with OCD would show increased oscillatory power in delta and theta frequency bands. Additionally, exploratory hypotheses were that there would be between-group differences in functional connectivity and 1/f parameters. Although several previous studies have investigated power spectral differences between OCD and HC groups, this is the first to account for the 1/f non-oscillatory activity and to use a cluster-based permutation method to account for multiple comparisons. Furthermore, this is the first study to compare and report findings of 1/f parameters between OCD and HC groups. The FC analysis was performed in adherence to a recently published checklist [23] designed to standardize connectivity analyses and to our knowledge, this is the first OCD FC study to conform to these standards.

## 2. METHODS

### 2.1. Participants

Male and female participants aged between 18 and 65 years were recruited through internet and poster advertisements and doctor referrals from the state of Victoria, Australia. The required sample size was calculated using Cochran’s formula (supplemental materials S1) [24].The clinical trial was conducted as per the latest version of the Guidelines for Good Clinical Practice [25]. Following verbal and written explanation of the nature of all involved procedures, informed written consent was obtained from each participant. All participants were reimbursed for their participation in the trial. The study received ethics approval from Monash Health Human Research Ethics Committee and was registered on the Australian New Zealand Clinical Trials Registry (ANZCTR).

The OCD group comprised individuals diagnosed with OCD according to the International Classification of Diseases – 10^th^ revision [World Health 26] or DSM-IV/V [American Psychiatric 27]. Symptom severity was assessed utilizing the Yale-Brown Obsessive Compulsive Scale (YBOCS) [28]. Additionally, the Beck Anxiety Inventory (BAI) [29] was used to assess the level of anxiety and the Quick Inventory for Depressive Symptoms – Self Report (QIDS-SR) [30] was used to assess the level of concurrent depression. Exclusion criteria for the OCD group included scoring <17 on the YBOCS, presence of an unstable medical/neurological disorder and being diagnosed with another mental health condition other than depression and anxiety. Participants were included regardless of their medication status but were required to be on a stable dose for at least 6 weeks prior to the study. Clinical data obtained from the patient group included age of OCD onset, illness duration, medication history, presence of comorbidities (depression and anxiety) and symptom severity.

Healthy control data was obtained from individuals who have never been diagnosed with a mental health or neurological disorder. Other exclusion criteria for the HC group included currently being on psychoactive medication and previous concussion or head injury with loss of consciousness for more than 10 minutes.

### 2.2. EEG Recording

EEG recording took place in a laboratory with constant levels of lighting and background noise from air conditioning. The participants were seated upright on a padded, comfortable chair and requested to close their eyes, try to stay awake and relaxed during the recording period. Prior to recording, participants were provided with an explanation of the procedure and its safety along with instructions to minimize muscle and eye movements that may affect the recording. All participants underwent 3 minutes of EEG recording at rest with eyes closed.

The EEG data was collected using the actiCHamp amplifier (Brain Products GmbH, Munich, Germany) with the BrainVision software (version 1.21.0303) and 64 Ag/AgCl electrodes embedded within an EasyCap (Herrsching, Germany) based on the international 10-20 system. Out of these, 63 electrodes were used (supplemental materials S2) and CPz was excluded due to being the reference electrode. All recordings were made at a sampling rate of 1000 Hz with the ground placed at AFz. A transparent electro-gel was injected onto the scalp to reduce impedance, which was kept below 5 kΩ. No online band-pass or notch filtering was applied during the recording.

### 2.3. EEG Pre-processing

Raw, continuous EEG data was pre-processed using the RELAX pipeline [31] implemented within MATLAB [32], which utilizes functions from EEGLAB [33] and fieldtrip [34]. This pipeline initially applied a 4^th^ order acausal Butterworth bandpass filter from 1 to 80 Hz and a second order acausal Butterworth notch filter from 47 to 53 Hz. Following this step, several measures were taken to identify and reject bad electrodes. The initial removal of noisy electrodes occurred through the “findNoisyChannels” function of the PREP EEG preprocessing pipeline [35]. Thereafter, electrodes were marked for rejection based on 1) extreme outlying amplitudes [36]; 2) extremely improbable voltage distributions; 3) extreme drift; 4) extreme kurtosis; and 5) muscle activity [37]. Overall, the rejection of a maximum of 20% of electrodes was accepted. If more than 20% were marked for removal, electrodes were ranked based on the number of epochs showing extreme artifacts and only the worst 20% were removed. The same measures were applied subsequently to mark extreme periods to be excluded from further analysis. The resultant data was then subjected to three sequential multiple Wiener filter (MWF) cleaning steps [38] to address the following artifacts: 1) Muscle activity: low-power log-frequency slopes of more than −0.59 were classified as epochs affected by muscle activity; 2) Blink artifacts: following bandpass filtering using a fourth order Butterworth filter from 1 to 25 Hz, pre-specified blink affected electrodes were selected and voltages were averaged within 1 s epochs. Blinks were flagged at time points where the averaged voltage exceeded the upper quartile plus thrice the inter quartile range of the distribution of all voltages in these pre-specified electrodes. The 800ms period surrounding each blink flag was marked as artifacts for further cleaning; 3) Horizontal eye movements and drift: Periods of selected lateral electrodes affected by horizontal eye movement showing voltages greater than twice the median absolute deviation (MAD) from the median of their overall amplitude, with a similar opposite voltage movement in the electrode on the opposite side of the head were classified as horizontal eye movements [39]. Epochs with an amplitude greater than 10 times MAD from the median of all electrodes were considered to be affected by drift [40] and marked as artifacts for further analysis. The resultant data was average rereferenced using the PREP method [35] before performing Independent Component Analysis (ICA) using fastICA [41]. Artifactual components were detected using ICLabel [42] and only these components were cleaned with wavelet enhanced ICA (wICA) [43]. Continuous data was then reconstructed back into the scalp space and previously rejected electrodes were spherically interpolated to obtain a full set of electrodes for each participant.

### 2.4. Detection of Better Oscillations with eBOSC

Following the initial pre-processing, data was further processed using an extended version (eBOSC) [44] of the Better OSCillation (BOSC) method [45]. This method was designed to detect the arhythmic, non-oscillatory portion of the EEG signal (1/f activity). In BOSC, duration (*D_τ_*) and power (*P_τ_*) thresholds are calculated by modelling the known background power spectrum. Thus, at a given frequency, BOSC detects increases in power above *P_τ_* of a specific minimum duration *D_τ_*, thereby removing non-repeating rises in spectral amplitude. In eBOSC, rhythm detection benchmarks are derived using simulations to further characterize rhythmicity. During the eBOSC process, 1s of data from bilateral edges were zero-padded to avoid edge artifacts and continuous data was segmented into 4s epochs with no overlap. The output of eBOSC includes the results of the time-frequency analysis as well as the specific time points of rhythmic episodes and this information was utilized in spectral analysis methods. Furthermore, the slope and intercept of the 1/f spectrum were also analyzed to identify any differences between HC and OCD groups.

### 2.5. Power Spectral Analysis

For spectral analysis, frequency bands were defined as: delta (0.5-4 Hz), theta (4-8 Hz), alpha (8-12 Hz), beta (12-30 Hz) and gamma (30-45 Hz). The wavelet time-frequency transformation table and the detected rhythmic episodes were obtained from the eBOSC output, and the power of each band was computed after removing the background 1/f activity. The output of periods showing oscillatory activity within each frequency band was separately time averaged for OCD and HC groups and cluster-based permutation was applied to the resultant data. The fieldtrip function “ft_prepare_neighbours” was used to find channel neighbors for spatial clustering using triangulation [34]. At least 2 neighboring electrodes were required to show significance to be included in a significant cluster. A family-wise error rate (FWER) was estimated by sampling and permuting the null distribution using the Monte Carlo method. A two-tailed independent sample t-test was used to evaluate the between-group differences. The cluster alpha (critical value used for thresholding the sample-specific t-statistics) was set at 0.05 and the number of permutations was set at 50000. Hypothesis testing was performed for delta, theta, and alpha bands, while beta and gamma bands were analyzed as exploratory analyses. The results of all primary hypothesis-driven analyses were controlled for experiment-wise multiple comparisons using a false discovery rate (FDR).

### 2.6. Functional Connectivity Analysis

FC analysis was performed adhering to a recently published EEG connectivity checklist that was developed to standardize FC studies while ensuring optimal connectivity assessment [23]. The debiased estimator of the weighted phase lag index (d-wPLI) and the coherence connectivity measures were separately used to compute the connectivity matrices based on the multi-taper Fourier spectral estimate. Both connectivity measures provide a value between 0 and 1 for each electrode pair, where higher values indicate higher connectivity.

Statistical analysis was performed using the Network-Based Statistic (NBS) which uses non-parametric permutation testing and controls for FWER using mass univariate testing at each connection [46]. The number of iterations performed for permutation testing was 10000 and the primary threshold was set at *p_t_* = 0.05. If a significant network was identified, the primary threshold was adjusted based on the recommendations of the NBS manual [46]; a range of thresholds were experimented where liberal thresholds (e.g., *p_t_* = 0.05) produced topologically extended results and conservative thresholds (e.g., *p_t_* = 0.001) produced strong, topologically focal results. The identified networks were visualized using the BrainNet Viewer [47].

As suggested in the checklist, continuous EEG data was segmented into 6s epochs, but the number of epochs was limited to the <50 category due to the short total duration of recording (the only non-optimal study parameter for d-wPLI). Re-referencing was performed with robust common average reference and all types of artefacts were addressed through the RELAX pre-processing pipeline. When our methods were applied to the checklist, a total score of 5 and 4.5 were achieved for d-wPLI and coherence measures (supplemental table 1), indicating high and moderate study quality, respectively.

### 2.7. Regression Analysis Between Symptom Severity and Oscillatory Power/Connectivity

A within-group linear regression was performed between the YBOCS and oscillatory power of each frequency band at each channel. A regression was also performed for each channel pair in any networks identified through the FC analysis. Bonferroni correction was used to account for multiple comparisons. These analyses provide information of any associations between the oscillatory power/connectivity measures and the OCD symptom severity.

## 3. RESULTS

### 3.1. Demographic and Clinical Data

The study sample comprised 25 OCD and 27 HC participants. The Demographic characteristics and clinical data are summarized in **Table 1**. No significant differences were observed between OCD and HC participants in any demographic variables.

**Table 1.** Demographic and Clinical Characteristics of Participants

| Variable | OCD (n = 25) |  |  | HC (n = 27) |  |  | Test statistic (*p*-val) |
| --- | --- | --- | --- | --- | --- | --- | --- |
|  | Mean | SD | n | Mean | SD | n |  |
| Demographic Characteristics |  |  |  |  |  |  |  |
| Gender (M/F) | | | 12/13 | | | 9/18 | $\chi^2 = 1.14, p = 0.29$ |
| Age (years) | 36.24 | 13.06 | 25 | 31.22 | 10.66 | 27 | $t = 1.52, p = 0.13$ |
| Handedness (R/L) | | | 21/4 | | | 26/1 | $\chi^2 = 2.22, p = 0.14$ |
| Marital status (S/M) | | | 20/5 | | | 21/6 | $\chi^2 = 0.04, p = 0.85$ |
| Clinical Characteristics |  |  |  |  |  |  |  |
| Age at onset (years) | 24.64 | 9.90 | 25 |  |  |  |  |
| Duration of illness (years) | 11.60 | 9.98 | 25 |  |  |  |  |
| YBOCS (total) | 28.00 | 3.82 | 25 |  |  |  |  |
| YBOCS - obsessions | 13.88 | 1.81 | 25 |  |  |  |  |
| YBOCS - compulsions | 14.12 | 2.39 | 25 |  |  |  |  |
| BAI | 17.04 | 8.92 | 24<sup>a</sup> |  |  |  |  |
| QIDS-SR | 10.17 | 4.94 | 24<sup>a</sup> |  |  |  |  |

*Note.* OCD – Obsessive Compulsive Disorder, HC – Healthy Control, SD – Standard Deviation, M – Male, F – Female, R – Right, L – Left, S – Single, M – Married, YBOCS – Yale-Brown Obsessive Compulsive Scale, BAI – Beck Anxiety Inventory, QIDS-SR – Quick Inventory of Depressive Symptoms-Self Report
<sup>a</sup> BAI and QIDS-SR scores for one participant were unavailable due to a data collection issue.

### 3.2. Power Spectral Analysis

Three clusters were identified through cluster-based permutation; two for the delta band (Cluster 1: *p* = 0.0065, cluster statistic = 84.81 ± 0.0007; Cluster 2: *p* = 0.046, cluster statistic = 26.38 ± 0.002) and one for theta (*p* = 0.041, cluster statistic = 44.46 ± 0.002). The topographical plots indicated that the OCD group had significantly higher power than HC in these frequency bands, in the regions highlighted by the clusters (**Figure 1**). Delta clusters were concentrated around the left fronto-temporal and the right parietal regions and the theta cluster was found at the left fronto-temporal and parietal regions. Upon controlling for multiple comparisons using FDR, only the first delta cluster was found to be significantly different (*p* = 0.0357). No significant clusters were found between OCD and HC groups in the alpha (*p* = 0.14, cluster statistic = 14.6 ± 0.003), beta and gamma bands.

**Figure 1.**
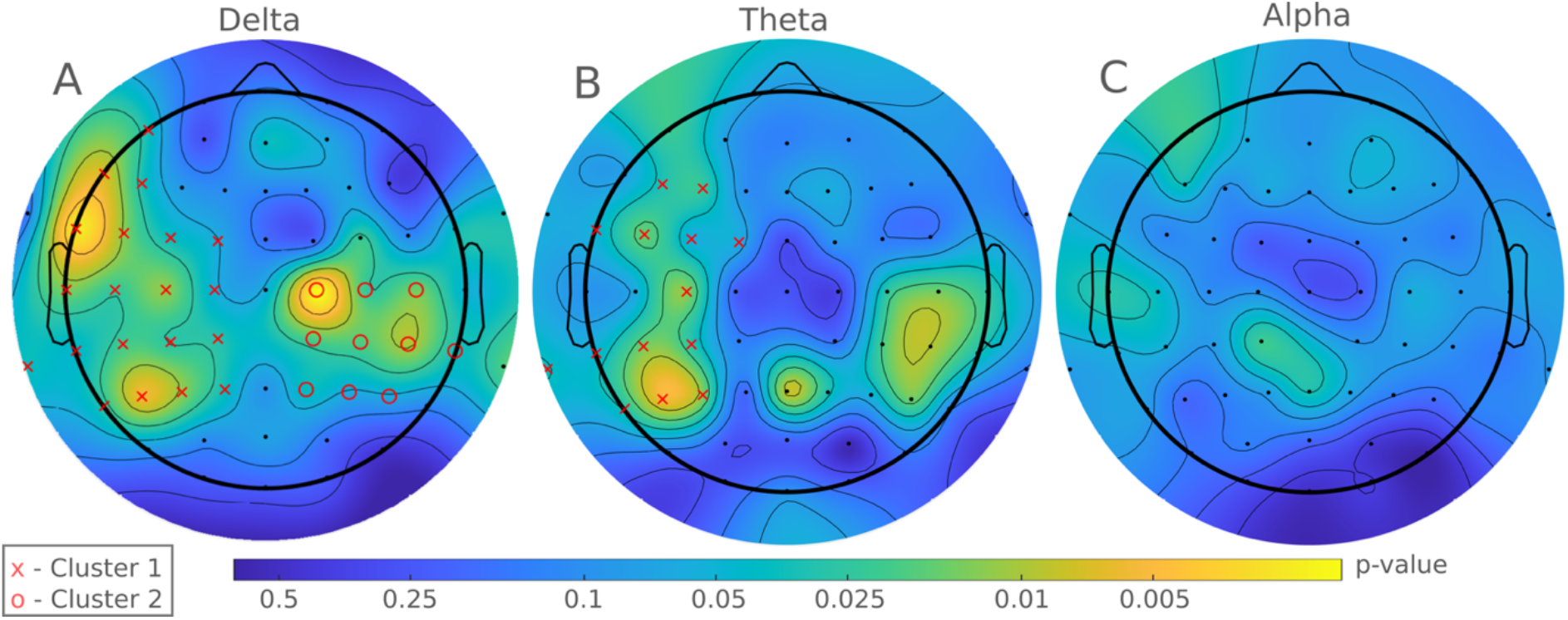
Topographical Plots Illustrating Spectral Power Differences Between OCD and HC Groups Using the Cluster-Based Permutation Method. *Note.* A) OCD groups showed significantly higher delta band power than HC in two significant clusters. Cluster 1 in the left fronto-temporal, parietal region (AF7, F5, F7, FC1, FC3, FC5, C1, C3, C5, T7, TP7, TP9, FT7, CP1, CP3, CP5, P1, P3, P5, P7), and Cluster 2 in the right parietal region (C2, C4, C6, CP2, CP4, CP6, P2, P4, P6, TP8. B) OCD groups showed significantly higher theta band power than HC in one cluster in the right fronto-temporal and parietal regions (F3, F5, FT7, FC1, FC3, FC5, TP7, TP9, P3, P5, P7, C3, CP3, CP5) C) No significant difference in band power was noted in the alpha band.

There were no significant differences between the OCD and HC groups for the intercept and slope parameters of the 1/f spectrum in any of the channels (supplemental figure 1).

### 3.3. Functional Connectivity

One significant network in the delta band was identified using the coherence measure (**Figure 2**). Following the adjustment of the primary threshold to the conservative value of *p_t_* = 0.0038, 21 nodes and 26 edges remained in the network (*p* = 0.049, network statistic = 3.13), indicating that the connectivity was significantly lower in the OCD group. Supplemental table 2 shows the unique connections identified within this network and the corresponding regions of interest. It is evident that multiple differences between groups in the strength of connections exist between the bilateral frontocentral and frontal electrode groups. No statistically significant differences in functional connectivity were identified between OCD and HC groups with the d-wPLI method. No other frequency bands had networks that were found to be significantly different between OCD and HC groups.

**Figure 2.**
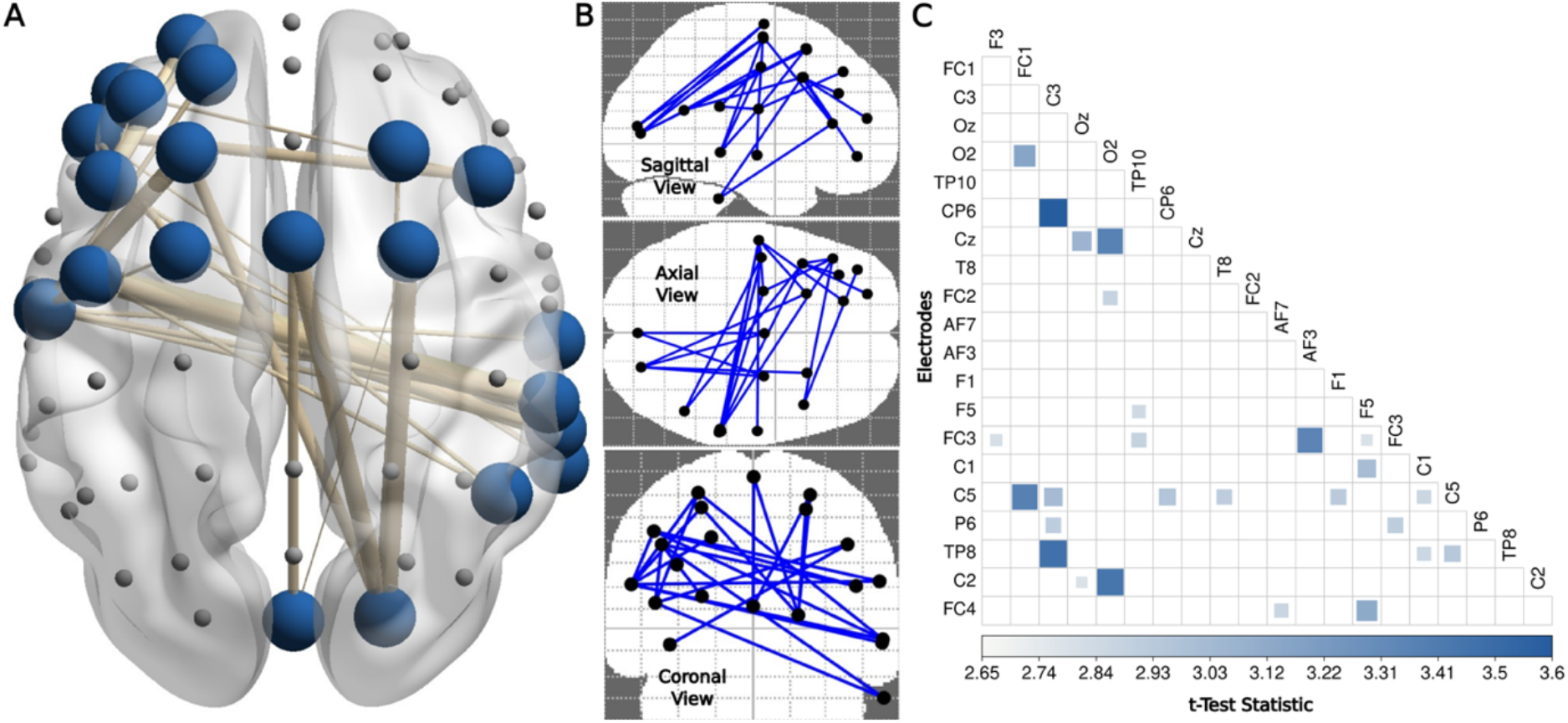
Network-Based Analysis of Functional Connectivity in OCD with Coherence Connectivity Measure. *Note.* A) Output of the BrainNet Viewer illustrating the identified connectivity network in the delta band. The nodes and weighted links indicate brain areas and the magnitude of the reduction in connectivity in the OCD group when compared to HC. B) Depiction of the connectome in three different views C) Connectivity matrix indicating the pairs of electrodes included in the identified network

### 3.4. Regression Analysis

Statistically significant associations were absent between the YBOCS and oscillatory power in all frequency bands following Bonferroni correction within the regression analysis. Furthermore, although a negative trend was identified between YBOCS and coherence connectivity (slope = −0.009, *p* = 0.057), the values were not statistically significant (supplemental figure 2).

## 4. DISCUSSION

EEG power spectral and functional connectivity findings in OCD populations have largely been inconsistent across previous studies. Therefore, we investigated spectral power changes between OCD and HC groups. Our study attempted to address these inconsistencies by adhering to better methodology involving a comprehensive EEG pre-processing pipeline and controlling for 1/f, non-oscillatory activity. Furthermore, the current study analyzed the parameters of the 1/f slope for the first time in an OCD sample. We also performed an exploratory analysis of network-based functional connectivity changes in OCD.

Power spectral findings of the current study showed that the OCD group had significantly higher power in the delta and theta bands in comparison to HC in several clusters of electrodes. The delta clusters were concentrated around the left and right fronto-temporal and parietal regions, while the theta cluster was in the left fronto-temporal region. These findings agree with several previous studies that reported increased delta and theta powers in these same regions in OCD [5–8, 11, 12, 19, 48–52]. However, two studies reported deficits in delta [53] and theta [54] power in OCD when compared to HC.

OCD is known to be associated with significant reductions in cerebral blood flow (CBF) in the fronto-temporal brain regions [55]. Insufficient rates of CBF, leading to poor cortical grey matter oxygen uptake, has been found to be correlated with increased slow EEG frequencies [56]. Additionally, as postulated by Michel et al., the increase in the power of slow frequency oscillations in the frontal regions could be due to over-activation of frontal slow wave generators causing excessive delta and theta powers [57]. Therefore, frontal slow wave generators being activated due to low frontal CBF may explain the raised slow wave oscillatory power in OCD. Furthermore, the presence of excessive frontal slow wave activity has also been reported in other mental health conditions including social phobia [58], depression [59] and schizophrenia [60]. Additionally, increased frontal delta activity has been reported in individuals with Alzheimer’s disease and mild cognitive impairment [61]. As depression is a common comorbidity of OCD [62], this finding may be an indicator of underlying depression, rather than OCD.

Although several previous studies reported increased [9, 10, 48, 52, 53] and decreased [11, 12, 50, 54, 63–65] alpha power in OCD groups when compared to HC, the current analysis did not find any differences in the alpha band. It has been reported that individuals with OCD who respond to SSRIs showed excess alpha power, while non-responders showed excess theta power [53]. The OCD sample recruited in this study comprised largely of non-responders to SSRIs, which might explain this finding.

There is limited literature available relating to EEG connectivity in OCD. The present study reported no differences in connectivity using the d-wPLI measure and significantly decreased functional connectivity using coherence, in a network comprising frontocentral, temporal and parietal nodes in the delta band. However, these results must be interpreted cautiously as coherence may find strong spuriously increased connectivity due to its vulnerability to volume conduction [66]. One previous study reported a similar observation, where global field synchronization in the delta band was found to be lower in OCD in comparison to HC [21]. Similarly, a resting-state magnetoencephalography (MEG) study reported decreased connectivity in the delta and theta bands in the OCD group when compared to HC [67]. Alterations of functional nature in OCD, such as neurotransmitter dysfunctions [68] and structural grey matter deformities [69] may explain the decreased connectivity findings of OCD. Literature suggests that synchronous brain oscillations in slow frequencies are crucial for cognitive and motivational processes of attention, memory, perception, planning and decision-making [70–72], all of which are known to be impaired in individuals with OCD. Therefore, impaired functional connectivity in the slow frequency bands may underpin the clinical symptoms of OCD.

The current study reported no significant differences in the 1/f parameters (slope and intercept) between OCD and HC groups, which potentially indicates that OCD is a condition that exhibits altered oscillatory power, but not alterations to 1/f parameters. Several previous studies have reported 1/f parameter changes in bipolar disorder [73], attention deficit hyperactive disorder [74] and age-related variations [75]. It has been reported that the 1/f slope and offset parameters are functionally relevant to excitation/inhibition balance [76] as well as neural firing strength/rate [77]. The lack of similar findings in our study may be due to OCD being a condition in which these are unaffected.

### 4.1. Limitations and Future Directions

The current study has several limitations. While the total sample size was sufficient to identify group-level differences as per the sample size calculation (supplementary S1), inclusion of additional participants would have increased the statistical power and therefore enabled sub-group comparisons to examine potential mediating effects of demographic or clinical variables which may influence EEG. For instance, controlling for mediating effects of medication status may be useful as the recruited OCD participants were not drug naïve, and were on different classes of medications which may influence EEG activity.

The resting EEG data was collected over a total duration of 3 minutes, which resulted in a smaller number of epochs than is optimal for connectivity analysis [23]. However, the extent of the influence of epoch number is not established, and the other parameters of our study met the optimal recommendations for connectivity analysis. A principal limitation of the d-wPLI method is its relative insensitivity to true connectivity findings when the coherency phase lies close to 0 or 180, phases at which coherence performs better at identifying true connectivity changes. However, coherence may produce false connectivity results due to its vulnerability to volume conduction [66]. Therefore, the findings of the current study should be interpreted cautiously. Furthermore, EEG connectivity studies for OCD are scarce in the literature and therefore, further connectivity analyses of OCD that adhere to the checklist and recommendations [23] are essential.

The regression analysis of the current study did not show a significant association between increasing symptom severity and reduced connectivity in the delta band. However, a negative trend was identified and the inadequate power may be due to the sample size insufficiency [78].

The current study is the first to incorporate the eBOSC pipeline in the EEG processing to account for 1/f non-oscillatory activity in an OCD study. Future EEG studies of OCD should also use a similar method in their analyses to validate these results. Due to the poor efficacy and the resultant non-adherence to medications, treating OCD with alternative novel modalities such as non-invasive brain stimulation is becoming popular [79]. The findings of the current study and other power spectral/connectivity studies may be used to determine parameters for future brain stimulation studies.

### 4.2. Conclusions

OCD is a mental health condition causing significant disability and the poor understanding of its pathophysiology has made it difficult to establish a definitive treatment. This study aimed to further the current knowledge of the disease by performing EEG power spectral and functional connectivity analyses. Our findings suggest that OCD is associated with raised oscillatory power in the delta and theta frequency ranges in fronto-temporal regions, while no differences were noted in the alpha band. The connectivity findings with the d-wPLI measure showed no differences between groups while a significantly reduced functional connectivity was found in the delta band using the coherence measure. These findings encourage future research to further explore oscillatory power and connectivity measures as potential biomarkers for OCD.

## ACKNOWLEDGMENTS

This work was supported by the Monash University Institute of Graduate Research (MPNP) and National Health and Medical Research Council of Australia Investigator grant 1193596 (PBF).

The authors assert that all procedures contributing to this work comply with the ethical standards of the relevant national and institutional committees on human experimentation and with the Helsinki Declaration of 1975, as revised in 2008. This study was approved by the Monash Health Human Research Ethics Committee (Reference: RES-20-0000-265A).

