## supplemental material for "Obsessive-Compulsive Disorder (OCD) is Associated with Increased Electroencephalographic (EEG) Delta and Theta Oscillatory Power but Reduced Delta Connectivity"

**Supplemental Materials**

S1. Sample size ($n_0$) was calculated using the Cochran's formula (Kotrlik & Higgins, 2001):

$$n_0 = \frac{Z^2 pq}{e^2},$$

where $e = 5.5\%$ denotes the margin of error, $p = 2\%$ is the estimated proportion of the population (Fullana et al., 2009), $q = (1 - p)$ and $Z = Z_{\alpha/2}$ is the standard normal probability of $\alpha = 0.95$ confidence level, which yields the required sample size of 25.

S2. EEG channels: FP1, Fz, F3, F7, FT9, FC5, FC1, C3, T7, TP9, CP5, CP1, Pz, P3, P7, O1, Oz, O2, P4, P8, TP10, CP6, CP2, Cz, C4, T8, FT10, FC6, FC2, F4, F8, FP2, AF7, AF3, AFz, F1, F5, FT7, FC3, C1, C5, TP7, CP3, P1, P5, PO7, PO3, POz, PO4, PO8, P6, P2, CP4, TP8, C6, C2, FC4, FT8, F6, AF8, AF4, F2, FCz

**Supplemental Table 1**

*EEG connectivity study quality checklist* (Miljevic, Bailey, Vila-Rodriguez, Herring, & Fitzgerald, 2021).

*Note.* Table 1a shows the checklist scoring for the analysis using weighted phase lag index (wPLI). Table 1b shows the checklist scoring for the analysis using coherence.

| Table 1a | 0 | 0.5 | 1 |
|---|---|---|---|
| 1. Re-referencing technique: | Single, mastoid / ear, nose, or not presented<br>☐ | CAR<br>☐ | REST, rCAR, or Laplacian<br>X |
| 2. Epoch length: | <3 s<br>☐ | 4-6 s<br>☐ | >6<br>X |
| 3. Number of sample epochs: | <50<br>X | 50-100<br>☐ | >100<br>☐ |
| 4. Artefact rejection technique (for specifics see "Checklist Scoring Specifics" section): | None<br>☐ | Noisy epochs, or channels rejected<br>☐ | All types of artefacts addressed<br>X |
| 5. Control for volume conduction (for specifics see "Checklist Scoring Specifics" section): | None<br>☐ | Lag, weighted, source, or Laplacian<br>X | (Lag or weighted) & (source-space or Laplacian)<br>☐ |
| 6a. Control for multiple comparisons (for specifics see "Checklist Scoring Specifics" section): | No post hoc<br>☐ | Invalid post hoc control or p-value = 0.01<br>☐ | Valid post hoc control (If network / cluster - based stats use 6b instead)<br>☐ |
| 6b. Only for studies assessing network metrics: | | Arbitrary thresholding / not model-driven<br>X | Model or data - driven: threshold or weighted<br>☐ |
| 7. Sample size estimation and consideration: | No considerations<br>☐ | Some consideration (i.e., *N* obtained from published literature)<br>☐ | Statistical consideration (i.e., a priori *N* calculation)<br>X |
| **Scores:** | ________ | 1.0 | 4.0 |
| **Total score & QR:** | 5.0 | | |

*Note.* 0 = not recommended for use; 0.5 = not optimal; and 1 = optimal for use.

| Table 1b | 0 | 0.5 | 1 |
|---|---|---|---|
| 1. Re-referencing technique: | Single, mastoid / ear, nose, or not presented ☐ | CAR ☐ | REST, rCAR, or Laplacian X |
| 2. Epoch length: | <3 s ☐ | 4-6 s ☐ | >6 X |
| 3. Number of sample epochs: | <50 X | 50-100 ☐ | >100 ☐ |
| 4. Artefact rejection technique (for specifics see "Checklist Scoring Specifics" section): | None ☐ | Noisy epochs, or channels rejected ☐ | All types of artefacts addressed X |
| 5. Control for volume conduction (for specifics see "Checklist Scoring Specifics" section): | None X | Lag, weighted, source, or Laplacian ☐ | (Lag or weighted) & (source-space or Laplacian) ☐ |
| 6a. Control for multiple comparisons (for specifics see "Checklist Scoring Specifics" section): | No post hoc ☐ | Invalid post hoc control or p-value = 0.01 ☐ | Valid post hoc control (If network / cluster - based stats use 6b instead) ☐ |
| 6b. Only for studies assessing network metrics: | | Arbitrary thresholding / not model-driven X | Model or data - driven: threshold or weighted ☐ |
| 7. Sample size estimation and consideration: | No considerations ☐ | Some consideration (i.e., *N* obtained from published literature) ☐ | Statistical consideration (i.e., a priori *N* calculation) X |
| **Scores:** | ______ | 0.5 | 4.0 |
| **Total score & QR:** | 4.5 | | |

*Note.* 0 = not recommended for use; 0.5 = not optimal; and 1 = optimal for use.

**Supplemental Figure 1**

*Analysis of the slope and intercept parameters of the 1/f slope*

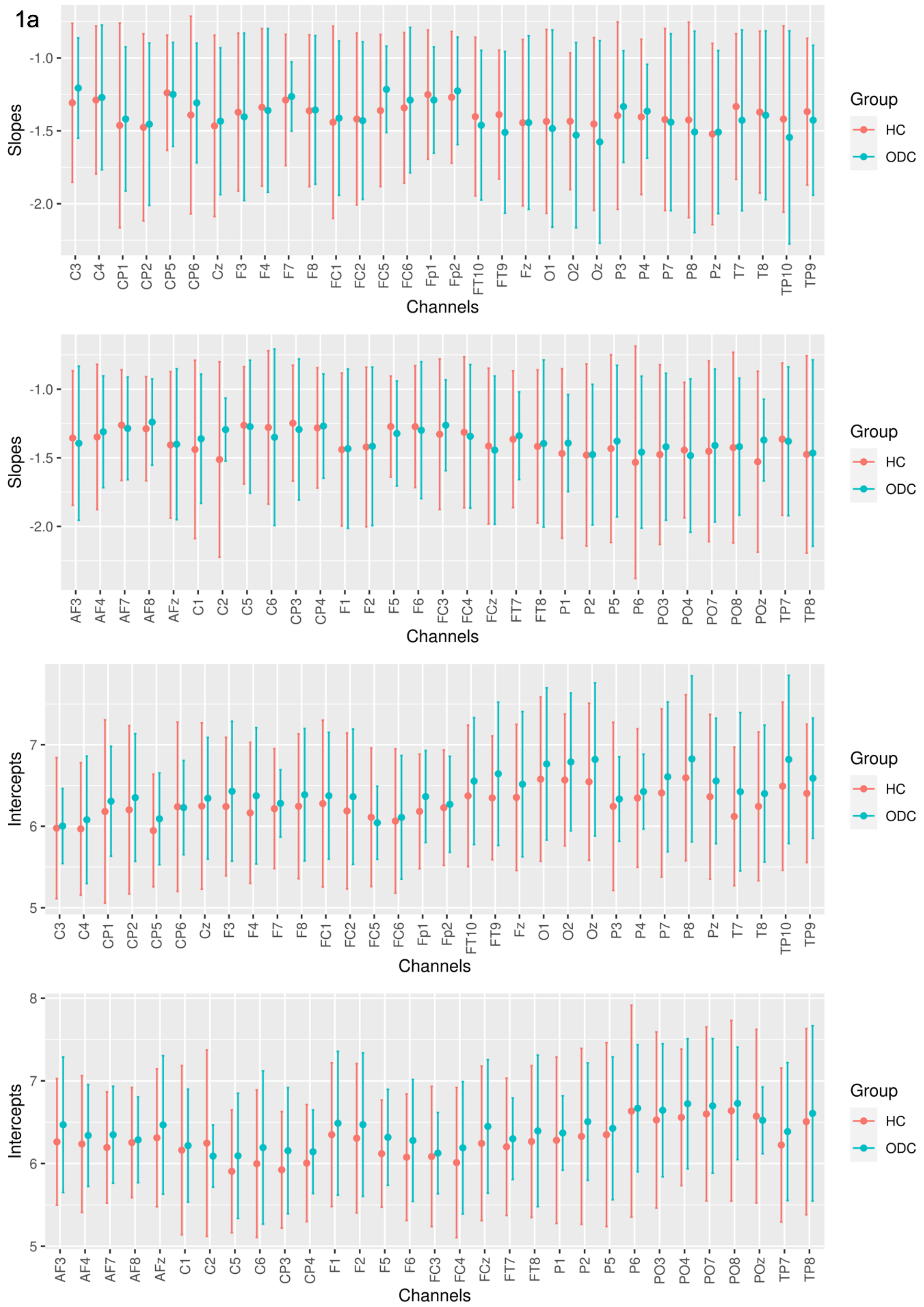

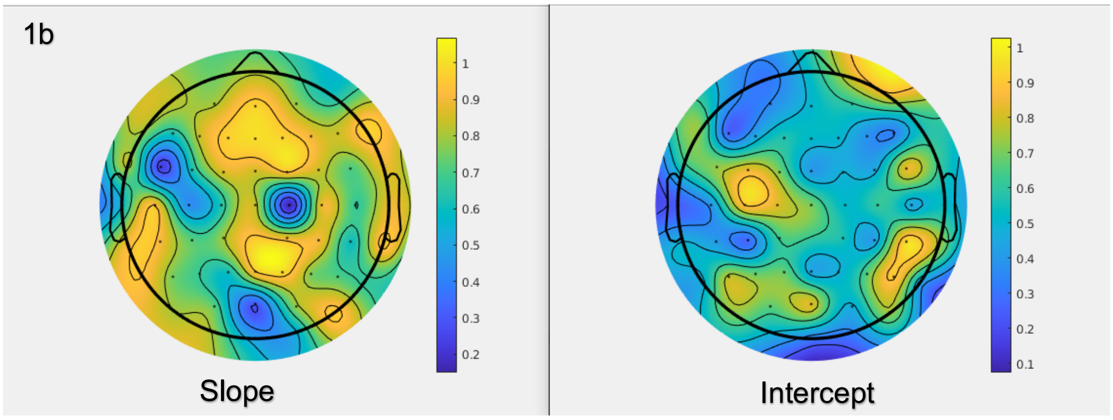


*Note.* The slope and intercept outputs from the eBOSC pipeline were analysed separately for OCD and HC groups. Figure 1a shows the mean values for each channel and figure 1b shows the topographical plots of the p-values of the t-test to compare the two groups. No significant differences were noted for either slope or intercept parameters between the OCD and HC group in any of the channels.

**Supplemental Table 2**

*Unique Connections of the Identified Delta Connectivity Network*

| Channels/nodes | Test statistic | Corresponding brain regions |
|---|---|---|
| C3 – CP6 | 3.60 | L-frontocentral – R-centroparietal |
| C3 – TP8 | 3.47 | L-frontocentral – R-temporal |
| O2 – C2 | 3.44 | R-occipitoparietal – R-frontocentral |
| O2 - Cz | 3.37 | R-occipitoparietal - Central |
| FC1 – C5 | 3.37 | L-frontocentral – L-frontocentral |
| AF3 – FC3 | 3.35 | L-frontal – L-frontocentral |
| FC1 – O2 | 3.16 | L-frontocentral – R-occipitoparietal |
| F5 – FC4 | 3.12 | L-frontal – R-frontocentral |
| Oz - Cz | 3.06 | Occipital - Central |
| C3 – C5 | 3.02 | L-frontocentral – L-frontocentral |
| F5 – C1 | 3.01 | L-frontal – L-frontocentral |
| CP6 – C5 | 2.96 | R-centroparietal – L-frontocentral |
| C5 – TP8 | 2.94 | L-frontocentral – R-temporal |
| F1 – C5 | 2.93 | L-frontal – L-frontocentral |
| T8 – C5 | 2.91 | R-temporal – L-frontocentral |
| C3 – P6 | 2.91 | L-frontocentral – R-centroparietal |
| FC3 – P6 | 2.91 | L-frontocentral – R-centroparietal |
| O2 – FC2 | 2.88 | R-occipitoparietal – R-frontocentral |
| TP10 – FC3 | 2.88 | R-temporal – L-frontocentral |
| C1 – C5 | 2.86 | L-frontocentral – L-frontocentral |
| AF7 – FC4 | 2.86 | L-frontal – R-frontocentral |
| C1 – TP8 | 2.85 | L-frontocentral – R-temporal |
| TP10 – F5 | 2.84 | R-temporal – L-frontal |
| F3 – FC3 | 2.81 | L-frontal – L-frontocentral |
| F5 – FC3 | 2.79 | L-frontal – L-frontocentral |
| Oz – C2 | 2.79 | Occipital – R-frontocentral |

*Note.* Nodes, weights of edges and brain regions of the delta band network identified through network-based analysis using the coherence connectivity measure

**Supplemental Figure 2**

*Regression Analysis Between Oscillatory Power/Connectivity and OCD Symptom Severity*

2a

| Frequency Band | $\beta$ (slope) | Standard Error | $t$ statistic | $p$-value |
|---|---|---|---|---|
| Delta | 0.254 | 0.167 | 1.514 | 0.144 |
| Theta | -0.316 | 0.147 | -2.145 | 0.043 |
| Alpha | -0.563 | 0.286 | -1.965 | 0.062 |
| Beta | 0.033 | 0.025 | 1.311 | 0.203 |
| Gamma | 0.005 | 0.002 | 2.211 | 0.037 |

2b

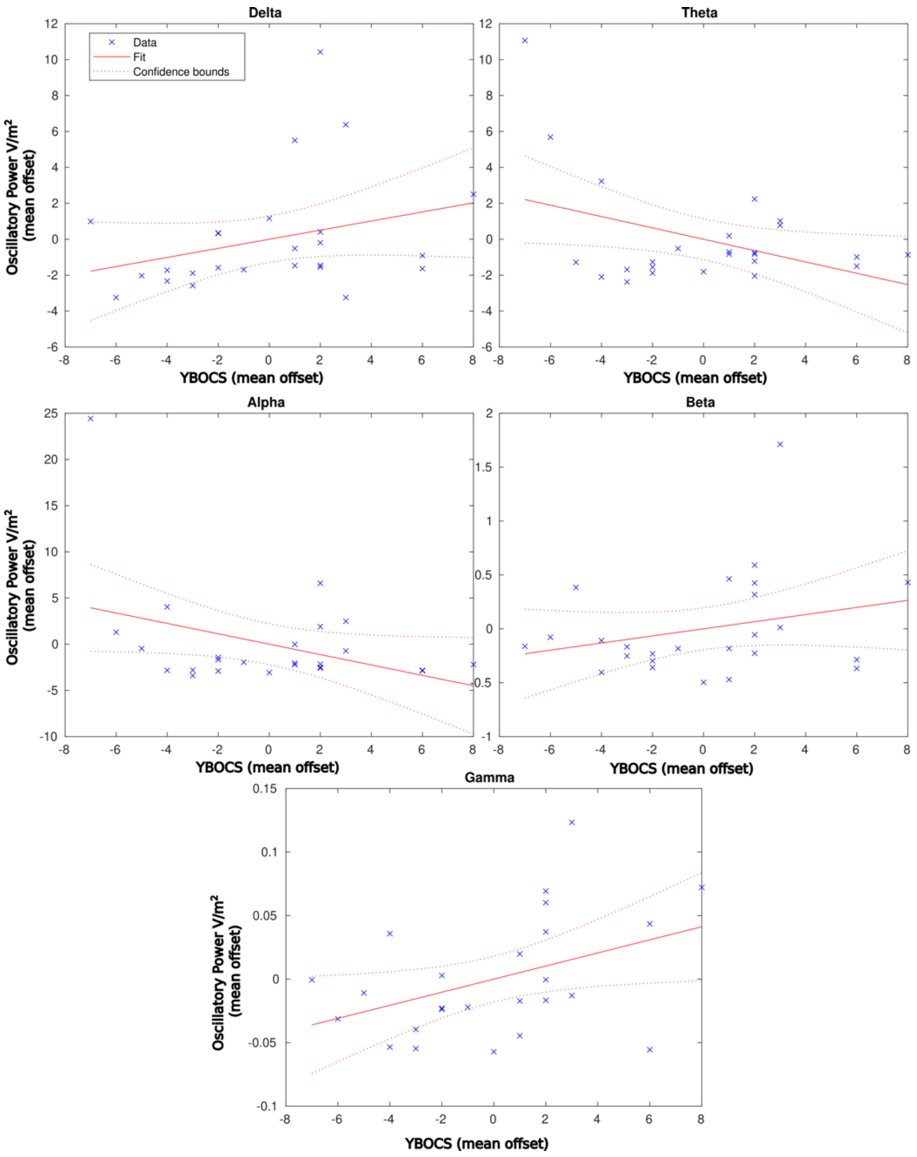


2c

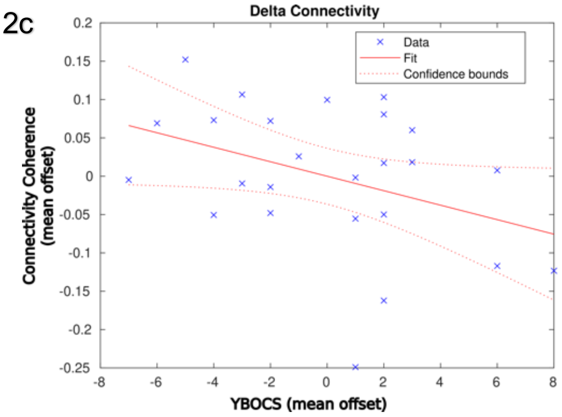


2d

| | |
|---|---|
| **$\beta$ (slope)** | -0.009 |
| **Standard Error** | 0.005 |
| **$t$Stat** | -2.005 |
| **$p$-value** | 0.057 |

*Note.* Figure 2a and 2b show the statistics and the scatter plot between the YBOCS (mean offset) and Oscillatory power (mean offset) for each frequency band at the electrode which showed the highest slope. Figure 2c and 2d show the same for coherence connectivity measure.
